# Functional dissociation of theta oscillations in the frontal and visual cortices and their long-range network during sustained attention

**DOI:** 10.1101/684829

**Authors:** Hio-Been Han, Ka Eun Lee, Jee Hyun Choi

## Abstract

Theta-band (4–12 Hz) activities in the frontal cortex have been thought to be a key mechanism of sustained attention and goal-related behaviors, forming a phase-coherent network with task-related sensory cortices for integrated neuronal ensembles. However, recent visual task studies found that selective attention attenuates stimulus-related theta power in the visual cortex, suggesting a functional dissociation of cortical theta oscillations. To investigate this contradictory behavior of cortical theta, a visual Go/No-Go task was performed with electroencephalogram recording in mice. During the No-Go period, transient theta oscillations were observed in both the frontal and visual cortices, but theta oscillations of the two areas were prominent in different trial epochs. By separating trial epochs based on subjects’ short-term performance, we found that frontal theta was prominent in good-performance epochs, while visual theta was prominent in bad-performance epochs, exhibiting a functional dissociation of cortical theta rhythms. Furthermore, the two theta rhythms also showed a heterogeneous pattern of phase-amplitude coupling with fast oscillations, reflecting their distinct architecture in underlying neuronal circuitry. Interestingly, in good-performance epochs, where visual theta was relatively weak, stronger fronto-visual long-range synchrony and shorter posterior-to-anterior temporal delay were found. These findings highlight a previously overlooked aspect of long-range synchrony between distinct oscillatory entities in the cerebral cortex and provide empirical evidence of a functional dissociation of cortical theta rhythms.

**IN BRIEF:** Previous literature emphasized the pro-cognitive role of coherent oscillatory networks between distal brain regions, such as the fronto-visual theta synchrony. However, such a conceptual framework has been challenged as recent findings revealed distinct behavioral correlates of theta oscillations found in different cortical regions, especially in the frontal and visual cortices. Here, we show that frontal and visual theta represent distinct cortical processes and that the functional connectivity between them increases during sustained attention, especially when one of the two theta rhythms is relatively suppressed. The data presented here highlight a novel aspect of neural long-range synchrony between distinct cortical oscillators with distinct functional significance in task performance.

## INTRODUCTION

In the mammalian brain, oscillatory networks in the lower frequency bands (< 12 Hz) such as the theta band (4–12 Hz) are often thought to be a key mechanism of brain-wide interaction between distal regions (von Stein and Sarnthein, 2000). In particular, increased theta-rhythmic activity of the frontal cortex is a hallmark of improved task performance and sustained attention, known to reflect the level of cognitive demand of the given task (Cavanagh and Frank, 2014; Clayton et al., 2015). In addition, theta oscillations of the frontal cortex have been known to form a phase-coherent network with task-related brain regions, including the sensory cortices (Benchenane et al., 2011). The emergence of a theta oscillation network reflects active neuronal processing for cognitive functions such as sustained attention (Clayton et al., 2015), cognitive control (Cavanagh and Frank, 2014), and working memory (Sauseng et al., 2010). For instance, Liebe *et al.* (Liebe et al., 2012) reported local-field potentials and unit activities that were synchronous across the prefrontal and visual areas in the theta band during visual working memory tasks.

On the other hand, recent findings suggested that theta activities of the visual area are distinct from those of the frontal cortex and that visual theta is also associated with attention but in an opposite manner: stimulus-evoked theta power decreased with increased attention. For instance, Spyropoulos *et al.* (Spyropoulos et al., 2018) recently reported that theta oscillations in the visual cortex elicited by visual stimuli decreased in amplitude when visual attention was directed. In addition, another recent study found arousal-reduced theta power in the high-order visual area (i.e., posterior parietal cortex) of the ferret brain (Stitt et al., 2018) despite its enhanced long-range theta synchrony with the prefrontal cortex during sustained attention reported by an earlier study (Sellers et al., 2016). In fact, a pioneering neuro-feedback study in the 1970s reported similar results of an association between occipital theta suppression and improved performance in a visual detection task (Beatty et al., 1974). There have also been several reports of enhanced theta in the posterior brain region being predictive of inattentiveness such as after sleep deprivation (Vyazovskiy and Tobler, 2005) or an attentional lapse (Mazaheri et al., 2009). Although the term “attention” of each study referred to different aspects of attention (e.g., internal or external attention (Chun et al., 2011)) on different time scales (e.g., minute or second levels), evidence generally suggests that the employment of attention yields contrary patterns of theta oscillations in the frontal and visual cortices.

This attentional modulation of frontal and visual theta in opposite directions poses a question about the functionality of theta oscillations in different brain regions, especially in terms of their long-range functional connectivity. If cortical theta oscillations of different regions have different roles with opposing behavioral correlates, how could they form a coherent oscillatory network? A possible explanation of this counterintuitive interaction between cortical oscillators is that the oscillations of one region are suppressed to achieve synchronization with those of the other region. Here, we hypothesize that enhanced fronto-visual theta synchrony can occur during the suppression of frontal or visual theta.

To address this hypothesis, we designed a forced-paced (i.e., the opposite of self-paced) visual Go/No-Go task for head-fixed mice and analyzed the electroencephalogram (EEG) oscillations of each area. With this design, we aimed to capture the dynamic interaction between frontal and visual theta within a single protocol. In particular, in No-Go trials, we expected the recruitment of both frontal and visual theta oscillations, depending on the attentive state of the animal. This is because elaborate sustained attention (i.e., remaining alert while inhibiting a Go response) has been known to elicit control-related theta in the frontal cortex (Clayton et al., 2015; Sellers et al., 2016), while visual perception without attention has been reported to elicit theta oscillations in the visual area (Spyropoulos et al., 2018). By tracking short-term behavioral performance within a single experimental session, we investigated state-dependent changes of frontal and visual theta oscillations and their coupling with higher frequency oscillations (e.g., gamma oscillations) to confirm their functional dissociation. Then, we characterized the property of functional connectivity between frontal and visual theta to show how distinct cortical oscillators are coupled to form a coherent long-range network.

## RESULTS

### Changes of behavioral performance during Go/No-Go task

We trained mice (n = 5) to associate rightward motion (S+) with reward (water) and downward motion (S-) with punishment (air puff) (Figs. 1A–B). The movement of visual stimulus ceased (i.e., black blank screen) at licking, as exemplified in Fig. 1C. After training (see Materials and Methods for details), we monitored the performance of the mice during the Go/No-Go test sessions for 10 successive days. During the test days, overall discrimination performance measured by d-prime (M ± SE, *d’ =* 0.475 ± 0.08) remained above the chance level (i.e., *d’* = 0, Wilcoxon signed-rank test, *Z* = 5.845, *p* < .001, see Extended Data Fig. 1-1 for details). There was no statistically significant difference in the discrimination performance across the test days (Kruskal-Wallis test, *χ*^2^ = 1.179, *p* = .999). Moreover, a paired comparison between performance of the first half (1–5 d) versus the last half (6–10 d) did not yield any statistically significant difference (Wilcoxon signed-rank test, *Z* = 0.405, *p* = .686), indicating the absence of a learning effect (>Fig. 1-1A).

**Figure 1.**
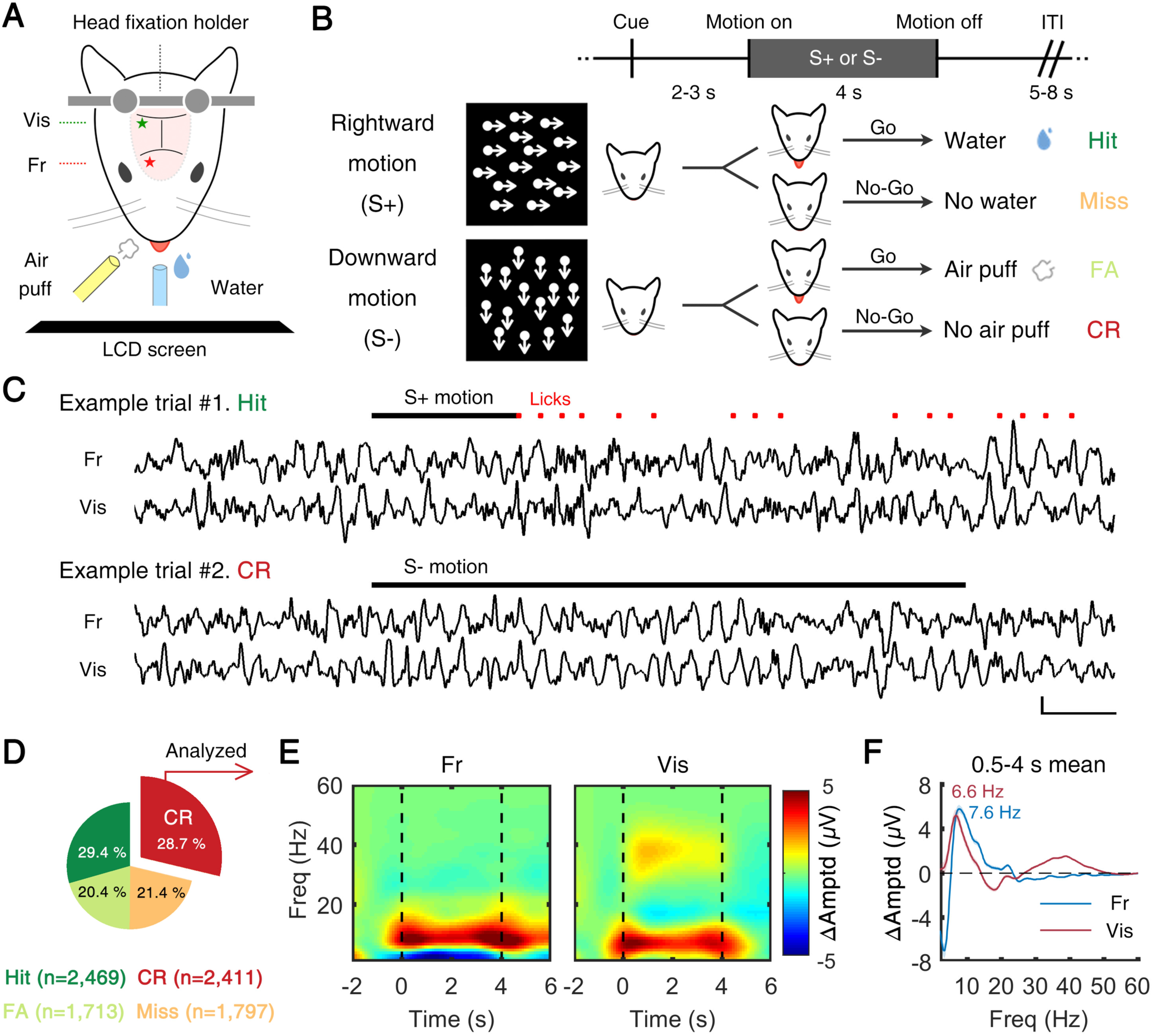
Theta oscillations during Go/No-Go task in frontal and visual cortice. (**A**) Schematic illustration of experimental setup. Mice performed Go/No-Go task in custom-built head restrainer. (**B**) Experimental procedures. Task required subject to discriminate direction of random-dot motion (rightward: S+, downward: S-). “Go” response (i.e., licking) to S+ and S-motion resulted in reward and punishment, respectively. There was no particular reinforcing stimulus following “No-Go” response (i.e., no licking). (**C**) Example EEG data of two trials. (Upper) For S+ motion, licking response was counted as “Hit,” and water reward was followed instantaneously. Motion stimulus was ceased at first lick (RT). (Lower) For S-motion, No-Go response was counted as “Correct rejection (CR).” Scale bars on bottom right indicate 500 ms and 300 µV. (**D**) Summary of behavioral results. From all subjects (five mice), and all sessions (10 days), data of 8,390 trials were collected. In this study, only the trials from CR (n = 2,411) were analyzed. (**E** and **F**) (**E**) Time-frequency representation of grand-averaged event-related spectral perturbation (ERSP) of each cortical area. In both the frontal and visual cortices, prominent theta (4–12 Hz) oscillations were observed during 4 s of stimulus presentation (No-Go period). (**F**) Temporal average (0.5 to 4 s) of theta oscillations in each cortical area. Numbers denote peak frequency of each channel. FA: false alarm, Fr: frontal cortex, Vis: visual cortex, Amptd: amplitude, Freq: frequency.

On the other hand, the animals showed clear changes in performance within a single session, probably due to the change of motivational factors (e.g., fatigue and quenched thirst) (Fig. 1-1B– C). The time-dependent patterns of performance were analyzed by dividing the session into early, middle, and late phases. When we classified the trials by Hit (licking to S+), CR (correct rejection; no licking to S-), Miss (no licking to S+), and FA (false alarm; licking to S-) (see Fig. 1D for the ratios and numbers of classified trials), the rate of correct answers did not show any time dependency: sum of Hit and CR ratios was 57.1%, 59.9%, and 57.4% in the early, middle, and late phases, respectively. Instead, the rates of licking decreased over time, shown through the sum of Hit and FA ratios which was 64.7%, 51.6%, and 37.1% in the early, middle, and late phases, respectively, possibly due to the decreased motivation for the task (see Fig. 1-1 for details).

### Prominent theta rhythms in frontal and visual cortices during correct rejection

All analyses of EEG data were focused on the time period from 0.5 s after stimulus onset until the cessation of stimulus (4 s after stimulus onset) during CR trials (n = 2,411 from five mice). As expected, our results showed that during the stimulus period of CR trials, theta-band activities increased in both the frontal and visual cortices (Figs. 1E–F). Elicitation of gamma activities in the low gamma band (around 40 Hz) was also observed in the visual cortex (Fig. 1E–F). The peak frequency of induced theta amplitudes was 7.62 Hz (FWHM = 6.26 Hz) and 6.64 Hz (FWHM = 4.96 Hz) in the frontal and visual cortices, respectively. The stimulus-locked and response-locked spectrograms in all trials are summarized in Fig. 1-2. Interestingly, although the behavior was the same (i.e., absence of licking response), Miss trials did not display such prominent theta band activity (Fig. 1-2).

### Assessment of attentional state using short-term task performance

Because of the nature of the Go/No-Go task, some trial epochs without active engagement in the task (e.g., attentional lapse) could be counted as correct No-Go responses. To distinguish those *by-accident* CR trials from the intended, *elaborate* No-Go responses, we tracked the changes in attentional state within a single session. During a task, attentional states change and correspondingly, performance fluctuates over time (Clayton et al., 2015). In human and non-human primate studies, the short-term history of correctness (Liebe et al., 2012) or reaction time (RT) (Esterman et al., 2013) is often used to track the change of behavioral state. Similarly, we tracked the short-term proportion correct, *p*_*c*_, by defining it as the ratio of correct trials (i.e., Hit and CR) across the 10 previous trials (see Materials and Methods for details). With a rough range of 2–5 minutes, *p*_*c*_ monitors the momentarily fluctuating, varying trial-to-trial, attentional state. The distribution of *p*_*c*_ for all CR trials exhibited a positively shifted Gaussian function (Fig. 2A, *μ* = .66, *σ* = .23, R^2^= .98) whose mean was above the chance level (i.e., *μ* = .5) (one-sample Z test, *Z* = 23.532, *p* < .001). Using the chance level as a threshold, we regarded the trial epochs as either attentive (i.e., *p*_*c*_ above the chance level; *p*_*c*_ > 0.5) or inattentive states (i.e., *p*_*c*_ ≤ 0.5). For CR trials, the lowest value of *p*_*c*_ was 0.2 and the highest value was 1, and attentive states were observed twice as frequently as inattentive states (1,550 attentive states, 861 inattentive states). When we drew RT as a function of *p*_*c*_ for licking cases (i.e., Hit and FA), RT decreased in the case of Hit trials (from 1.93 s for *p*_*c*_ = 0.2 to 1.14 s for *p*_*c*_ = 1, *Z* = 3.047, *p* = .002, Fig. 2B). On the other hand, RT in the case of FA trials did not show any dependence on *p*_*c*_. In this way each trial was labeled with *p*_*c*_, which successfully reflected the change of attentional state, for further analysis.

**Figure 2.**
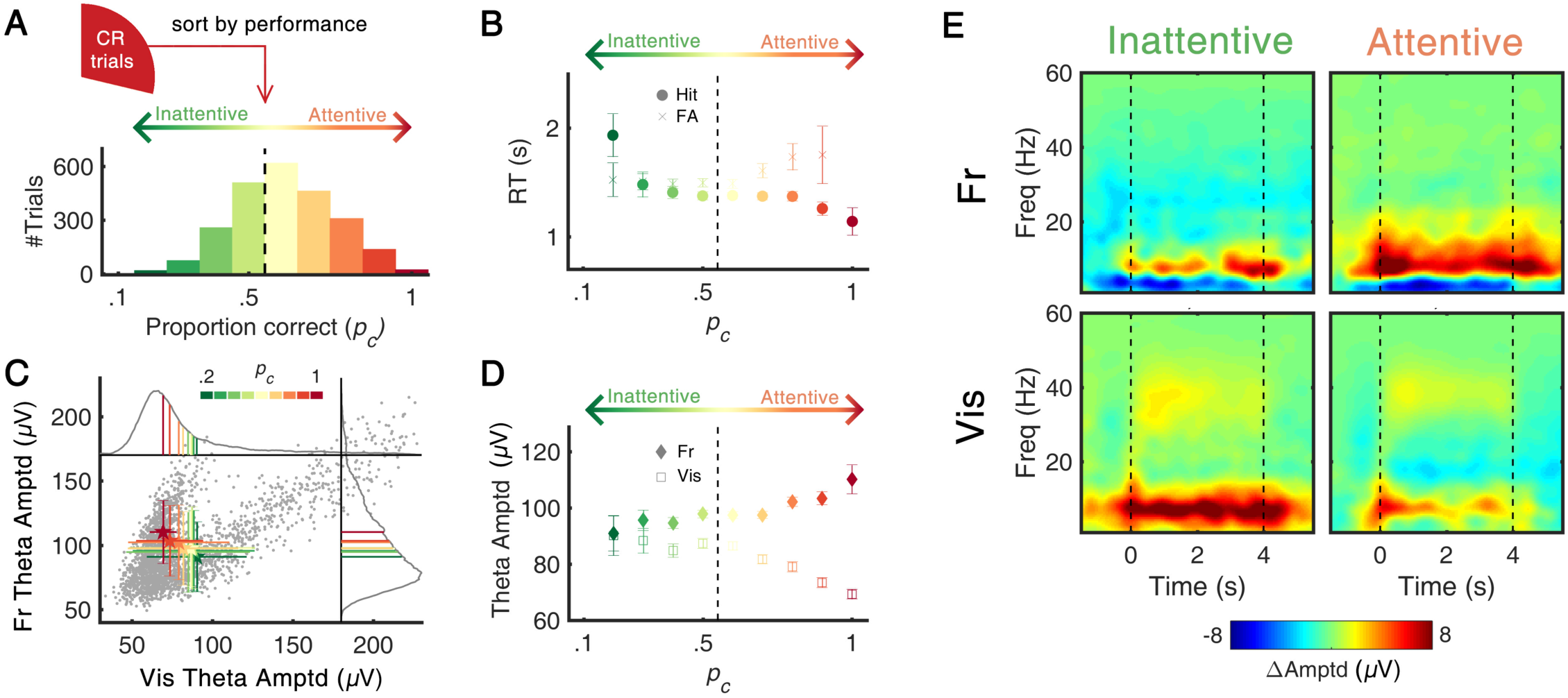
Task performance and theta amplitude. (**A**) Distribution of *p*_*c*_ (proportion correct) in all CR trials (n = 2,411). *p*_*c*_ was calculated across 10 previous trials. With a threshold of chance level (*p*_*c*_ = 0.5), each trial was classified as either attentive (*p*_*c*_ > 0.5) or inattentive (*p*_*c*_ ≤ 0.5) state. (**B**) Change of mean RT as function of *p*_*c*_. The RT of Hit trials decreased as *p*_*c*_ increased (Pearson’s *r* = -.051, *p* < .05), suggesting the change of attentional level within the Hit trial epochs. (**C**) Scatterplot of frontal/visual theta amplitude with color-coding of *p*_*c*_. Each gray dot represents the observed theta amplitude value of a single CR trial. Gray solid lines indicate the probability density function of each channel. Along with the changes of *p*_*c*_, the mean amplitudes of frontal and visual theta showed a negative relationship. (**D**) Mean amplitude of frontal/visual theta as function of *p*_*c*_. The linear correlation between theta amplitude and *p*_*c*_ was positive in the frontal (*r* = .076, *p* < .001) and negative in the visual (*r* = -.099, *p* < .001), suggesting their distinct behavioral correlates. (**E**) Time-frequency representation of oscillatory amplitudes highlighting state- and region-dependent change of theta amplitudes. Black dashed lines indicate the start/end of the visual stimuli. Error bars represent ±1 *SD* (**C**) and *SE* (**B, D**) of the means. *p*_*c*_: probability correct, RT: reaction time.

### Opposite effects of attentional state on frontal and visual theta amplitudes

The primary aim of the experiment was to articulate the ongoing theta activity signatures with respect to attention, which were reported to act in an opposite way in the frontal (Clayton et al., 2015) and visual (Spyropoulos et al., 2018) cortices. Thus, we were interested in the differences in ongoing theta activities between two attentional states. The scatterplot in Fig. 2C shows a positive correlation between frontal and visual theta amplitudes. On the other hand, the mean of each *p*_*c*_ presented as a colored error bar shows a negative correlation, with a movement in their distribution from the lower-right to the upper-left corner as *p*_*c*_ is increased. This opposite behavior of frontal and visual theta is manifested in the plot of theta amplitudes as a function of *p*_*c*_ (Fig. 2D). In the frontal cortex, the correlation between theta amplitude and *p*_*c*_ was positive, reaching its maximum at *p*_*c*_ = 1 (Pearson’s *r* = +.08, *p* < .001). In contrast, the relationship was negative for visual theta, reaching its minimum at *p*_*c*_ = 1 (Pearson’s *r* = -.10, *p* < .001). The average theta amplitudes for attentive versus inattentive, analyzed in the time-frequency domain, demonstrates the opposite effect of attention on the frontal and visual theta amplitudes (Fig. 2E). In the frontal cortex, theta amplitude in attentive states was significantly larger compared to that in inattentive states (96.5 ± 0.8 *μ*V in inattentive, 99.1 ± 0.7 *μ*V in attentive; Wilcoxon rank-sum test, *Z* = 3.189, *p* < .001). On the other hand, in the visual cortex, the amplitude of theta in attentive states was significantly smaller compared to that in inattentive states (86.7 ± 1.0 *μ*V in inattentive, 82.2 ± 0.9 *μ*V in attentive; Wilcoxon rank-sum test, *Z* = −2.582, *p* < .001). These contrasting correlations suggest that frontal and visual theta activities might have different functional roles during the engagement of cognitive processes required for the No-Go response, raising a question about the neurological signature of sustained attention.

### Distinct pattern of phase-amplitude coupling with fast oscillations

To further investigate the role of theta in each region, we performed the phase-amplitude cross-frequency coupling (CFC) analysis, whose characteristics are suggested to reflect the architecture and function of underlying neuronal circuits (Hyafil et al., 2015). The amplitude maps of fast oscillations plotted with respect to theta phase revealed strong but different CFC patterns across different *p*_*c*_ values and regions (Fig. 3A). For example, in the high-performance epochs with higher *p*_*c*_, the amplitude of fast oscillations was strongly modulated by the phase of theta oscillation in the frontal cortex compared to in the low-performance epochs. To analyze the effect of behavioral state on CFC, we calculated the modulation index (MI) (Tort et al., 2010) as an indicator of CFC strength. As depicted in Fig. 3B, the MI in the frontal cortex showed frequency-specific modulation of behavioral state, while the visual cortex did not show such modulation. Between theta and low gamma (20–40 Hz), the attentional enhancement in the frontal CFC was statistically significant (inattentive: 6.1 × 10^−4^ ± 0.1 × 10^−4^, attentive: 6.6 × 10^−4^ ± 0.1 × 10^−4^; Wilcoxon rank-sum test, *Z* = 2.187, *p* = .029) (Fig. 3C). For the faster frequency band (high gamma, 80–160 Hz) in the frontal cortex, attentional reduction was observed (inattentive: 7.4 × 10^−4^ ± 0.2 × 10^−4^, attentive: 7.0 × 10^−4^ ± 0.1 × 10^−4^; Wilcoxon rank-sum test, *Z* = −2.546, *p* = .011). However, in the visual cortex, the attentional effect on CFC was not statistically significant for either low gamma (*Z* = −1.443, *p* = .149) or high gamma (*Z* = 1.121, *p* = .262). It should be taken= into consideration that the CFC changed due to the theta amplitudes, rather than behavioral performance, as previous studies have both shown that CFC can be either dependent to amplitude of ongoing theta oscillations (Canolty et al., 2006; Tort et al., 2008) or independent to it (Kendrick et al., 2011). To elucidate possible confounding, an additional single-trial GLM (Generalized Linear Model) analysis was performed, and the results suggest that both factors – theta amplitude and attentional state – affects CFC (See Extended Data Figure 3-1).

**Figure 3.**
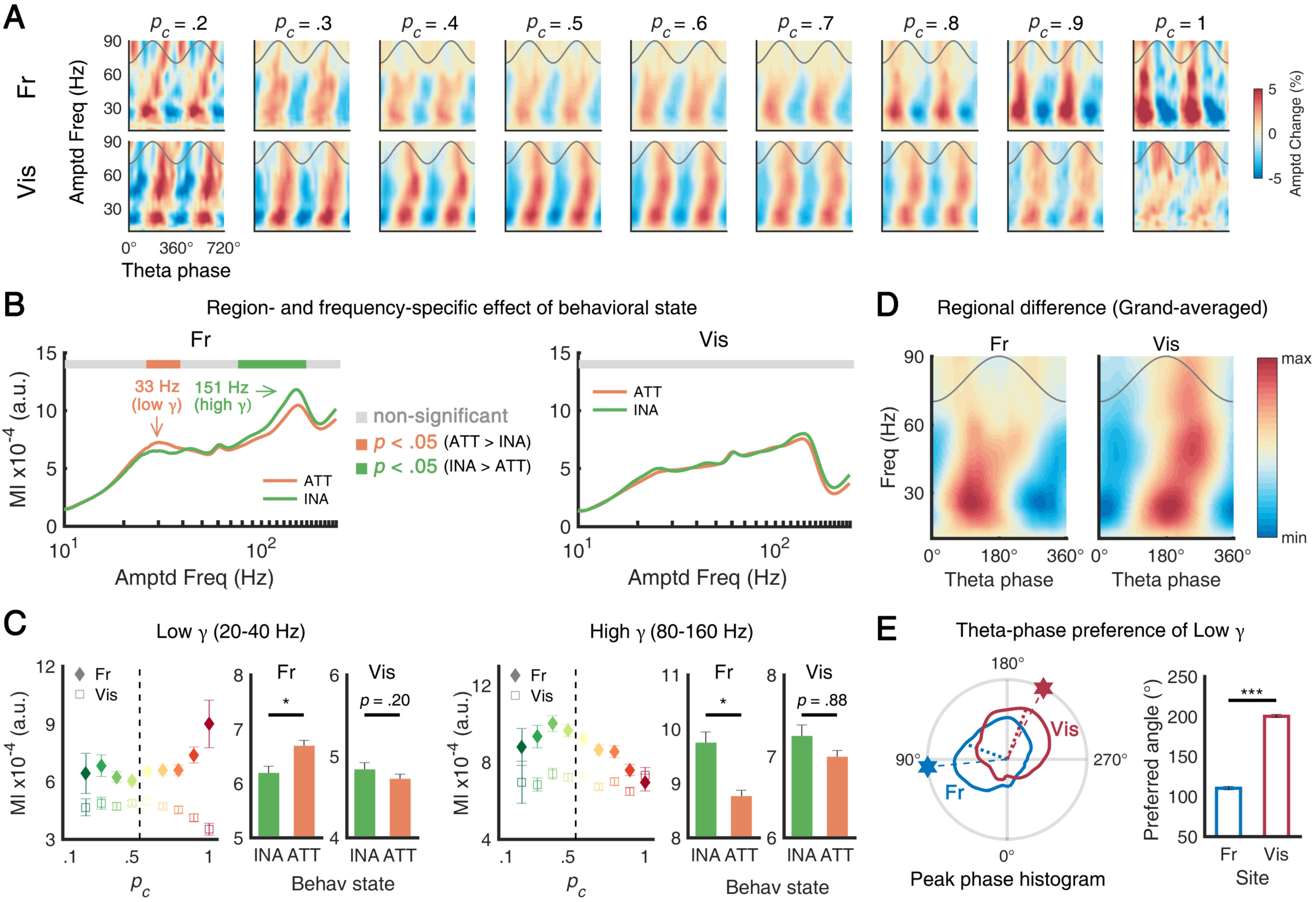
Task performance and theta-gamma phase-amplitude couplin. (**A**) Color-coded fast-frequency amplitudes as function of theta phase in frontal (top) and visual (bottom) and as function of *p*_*c*_ (left to right). (**B, C**) MI for local theta-gamma coupling as function of *p*_*c*_ (left: frontal, right: visual). In the frontal, the effect of attention on CFC was frequency-specific (increased CFC with low gamma, decreased CFC with high gamma). The attentional modulation of the visual coupling was not significant. (**D**) Grand-averaged amplitudes of fast oscillations as function of theta phase, showing noticeable regional difference of preferred theta phase (left: frontal, right: visual). (**E**) Trial histogram of preferred theta phase of 20–40 Hz oscillations (left). Hexagons and dotted lines denote the peak and mean of each polar histogram, respectively. Mean preferred angle of each area (right). The distances between preferred angles over sites were both significant. Error bars represent ±1 *SEM*. * *p* < .05, *** *p* < .001. CFC: cross-frequency coupling. INA: inattentive, ATT: attentive. MI: modulation index.

These region-specific behaviors of CFC were further analyzed by investigating the theta phase at which gamma amplitudes peaked (i.e., peak preference). This was done to check that the two theta oscillations are qualitatively distinct, as previous studies have shown that such peak phase difference implies differences in the underlying circuit motif (See Fig. 4 in (Voloh and Womelsdorf, 2016)). When theta phase angles were pooled separately for each region, their distributions differed (see polar plots in Fig. 3D). In the frontal cortex, the preferred theta phase (i.e., a phase bin of theta where maximum low gamma amplitude was detected) was observed in the middle of the rising phase (109.7 ± 2.0°), whereas in the visual cortex, the preferred theta phase was observed near the start of the falling phase (191.1 ± 1.7°). The distributions of the preferred theta phase of low gamma were statistically different for the frontal cortex and the visual cortex (circular Kruskal-Wallis test, χ^2^ = 251.140, *p* < .001). The angle distance of the preferred theta between them was 81.4°, corresponding to 32.3 ms at 7 Hz. In addition to the results from distinct behavioral modulation (Figs. 2C–E), the result of our CFC analysis presented here provides more reliable evidence that the theta rhythms of each area have distinct functional roles in the cortical microcircuit.

**Figure 4.**
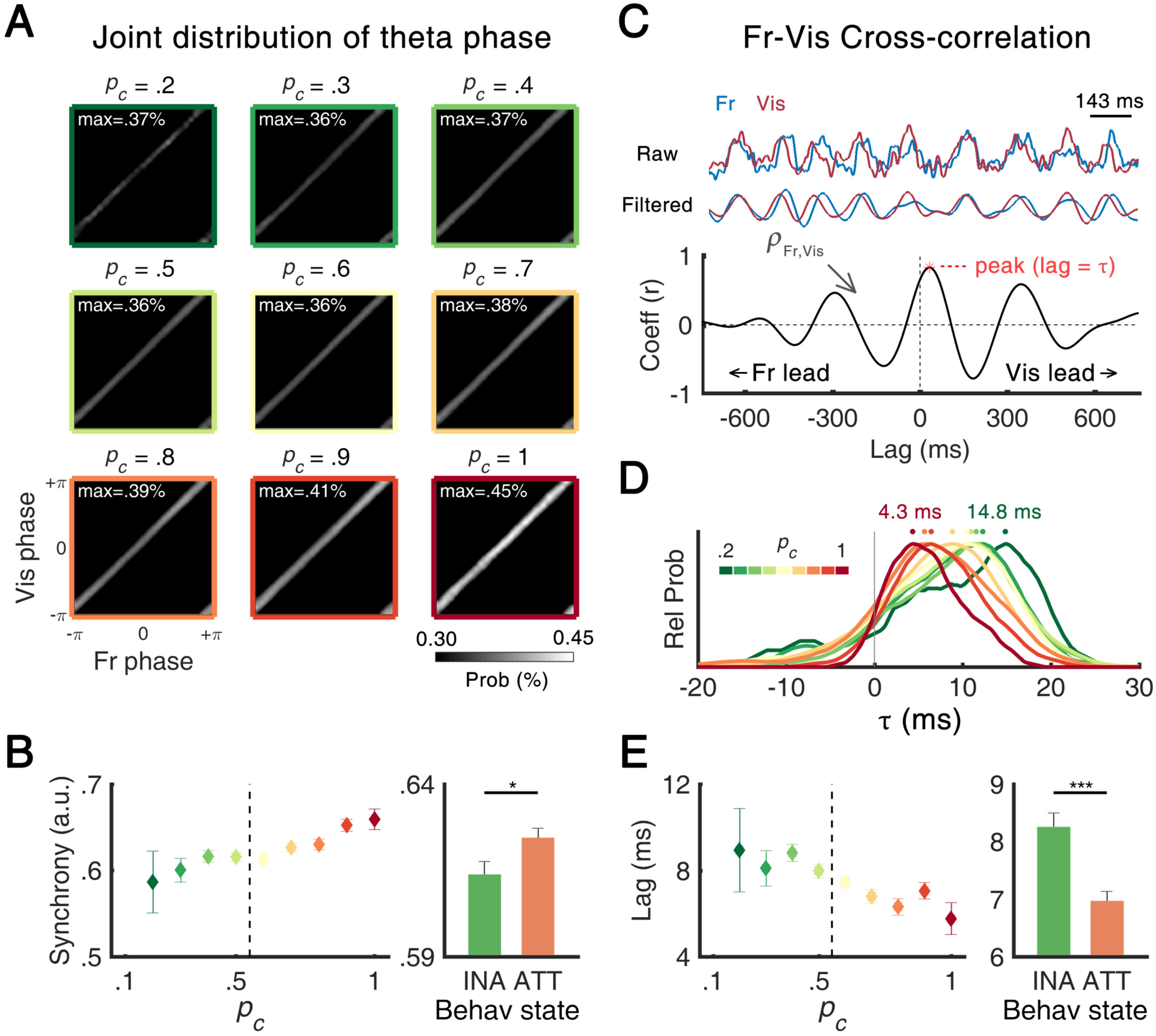
Synchrony and lags of theta oscillations. (**A**) Two-dimensional representation of joint phase histogram of frontal and visual theta oscillations as function of *p*_*c*_. The white color of each histogram indicates the higher probability of colocalization of the phase of two oscillations, suggesting a higher level of synchrony between two oscillations. The in-set text indicates the maximum value of colocalization probability in each *p*_*c*_ bin. (**B**) Mean fronto-visual phase synchrony calculated by PLV as function of *p*_*c*_. The phase synchrony was higher for the attentive state than the inattentive state. (**C**) Cross-correlation analysis of example EEG epoch. (Top) To calculate temporal lag (τ), raw time traces of EEG data from two channels were band-pass filtered in the theta band. (Bottom) After filtering, the cross-correlation function was obtained and τ was defined as a temporal lag where the correlation coefficient reaches its maximum. Positive and negative values of τ indicate a lead of the visual and frontal theta, respectively. (**D**) Trial histograms of τ between frontal and visual theta. The dots denote the peak of each histogram. Color-coded texts indicate the peak position from the lowest *p*_*c*_ (green) and to the highest *p*_*c*_ (red) bin. (**E**) Mean τ as function of *p*_*c*_. The τ was smaller in the attentive state than in the inattentive state. Error bars represent ± 1 *SEM*. * *p* < .05, *** *p* < .001. INA: inattentive, ATT: attentive, PLV: phase locking value.

### Enhanced fronto-visual functional connectivity during attentive state

Phase synchrony analysis allows the examination of signals that are cooperative between brain areas. To investigate the dynamic interplay of the fronto-visual theta network as attention fluctuates, we first examined the phase relationship between two theta rhythms using a joint phase histogram (see Materials and Methods for details). Fig. 4A highlights the systematic change of the instantaneous phase relationship as a function of *p*_*c*_. In general, the joint phase of frontal and visual theta showed the stronger colocalization along the diagonal axis regardless of behavioral state, which indicates their intimate phase relationship. Furthermore, the colocalization along the diagonal axis became more robust as *p*_*c*_ increased, suggesting a higher level of phase synchrony between the two theta rhythms. The phase locking value (PLV) (Lachaux et al., 1999), an amplitude-independent parameter, elucidates the brain state-dependent synchrony between frontal and visual theta oscillations (Fig. 4B). Compared to the inattentive state, the PLV was significantly higher in the attentive state (PLV = 0.614 ± .004 in inattentive, 0.624 ± .003 in attentive; Wilcoxon rank-sum test, *Z* = −2.313, *p* = .021). Our result of a fronto-visual theta synchrony associated with task performance is consistent with a previous finding in a non-human primate study (Liebe et al., 2012). While an opposite relationship between frontal and visual theta amplitudes was found in association with the behavioral state, a stronger fronto-visual long-range functional network was established, especially when frontal theta was relatively increased and when visual theta was relatively suppressed.

The asymmetrical joint phase distribution (i.e., bottom-right corner of each joint phase histogram in Fig. 4A) raised the possibility of a driver–responder relationship between frontal and visual theta. Therefore, we investigated the time lag, *τ*, of frontal theta with respect to visual theta by finding the peaks at the cross-correlation functions (CCFs) of the two theta rhythms. Fig. 3C illustrates an example CCF of frontal and visual theta rhythms. *τ* was obtained at each trial and the distributions of *τ* were plotted for different *p*_*c*_ values (Fig. 4D). Among all trials, a majority of the trials (84.61%) had positive *τ*, indicating that visual theta predominantly precedes frontal theta. In addition, as *p*_*c*_ increased, the distribution of *τ* shifted to shorter phase lag. The peak *τ* in Fig. 4D systematically decreased as the state changed from inattentive to attentive, with values of 14.83 and 4.33 ms at *p*_*c*_ of 0.2 and 1, respectively. *τ* was 8.25 ± 0.24 and 6.97 ± 0.17 ms for inattentive and attentive states, respectively (Wilcoxon rank-sum test, *Z* = 4.832, *p* <.001). In addition, we found that the peak frequency of the cross-spectral density function was significantly higher for the attentive state (5.84 ± 0.03 Hz) compared to the inattentive state (5.76 ± 0.04 Hz) (Wilcoxon rank-sum test, *Z* = 2.052, *p* = .040) (Fig. 2-1). The increase of synchrony and shift to shorter phase lag during the attentive state suggests the contribution of theta oscillations to task engagement and sustained attention via their long-range functional network.

It may seem odd that we calculated the phase synchrony after demonstrating that the two oscillators are distinct in peak frequency, since two oscillations must have transient periods of same frequency for phase locking to occur. Indeed, when we calculated synchrony for various instantaneous theta frequency, we found there were times during which the two theta rhythms oscillated in synch (See Extended Data Fig 4-1). Frequency difference was low when high phase synchrony was observed, and the probability of converging frequency difference (to 0) was found to be modulated by attention, which explains why PLV was higher during the attentive state.

## DISCUSSION

We found that the frontal and visual theta induced by the task had functionally distinct features. Whereas the frontal theta amplitude was positively correlated with behavioral performance, the visual theta amplitude was negatively correlated. In the frequency domain, the peaks of the two theta oscillations differed by almost 1 Hz, and their phases modulated fast cortical oscillations in different ways. Nonetheless, the frontal-visual theta synchrony increased as a function of performance despite the weakened amplitude of visual theta, implying a critical role of phase-locked oscillations in successful task engagement. It is unlikely that frontal and visual theta have common driving mechanisms; instead, it is more likely that the two theta have independent origins but are weakly coupled to each other. We should note that there is a distinction between phase locking (i.e., coupled oscillation with a constant phase difference) and phase trapping (i.e., un-locked oscillation with the same frequency) in nonlinear oscillator systems (Aronson et al., 1990). Indeed, the frontal and visual theta in our observation showed a phase transition from a stable phase-trapping system to a non-symmetric phase-locking system as the attention level increased. Although the differences in the values were small, we observed that there were abrupt changes in the interaction parameters (e.g., max in joint phase histogram, synchrony, and lag in Fig. 4) near *p*_*c*_ = 0.8. Furthermore, the behavioral parameters (i.e., RT in Fig. 2B) and the neural dynamics parameters (i.e., theta amplitude in Fig. 2D, theta–gamma coupling in Figs. 3A and 3C) presented their characteristic features near *p*_*c*_ = 0.8. It is highly possible that the theta oscillations changed their dynamic trajectory at the highly task-engaged brain state in their phase portrait; however, determining whether this transition contributed to sustained attention or vice versa was beyond the scope of this study.

Our phase dynamics analysis showed that visual theta led frontal theta throughout the recording sessions. At the same time, frontal, not visual, theta increased its amplitude at higher *p*_*c*_ accompanied by a reduction in its phase delay to visual theta. This confounding behavior at higher *p*_*c*_ could be an influence of top-down directed frontal theta for cognitive control such as motor inhibition (Cavanagh and Frank, 2014) or phase resetting of theta oscillations by a stimulus in a highly task-engaged brain state (Rizzuto et al., 2003). In terms of the *Communication through Coherence* theory (Fries, 2015), the tight interplay of two oscillations renders neuronal communication more effective, precise, and selective. It is possible that a new common source arose around *p*_*c*_ = 0.8 to enhance their synchrony and that the newly adjusted phase lag was simply differential conduction delay. However, the different patterns of theta–gamma coupling suggest this was not the case. Recently, Zhang *et al.* reported that human cortical oscillations were regionally distinctive and had the properties of weakly coupled oscillations rather than those of oscillations driven by a common source or feedback loop (Zhang et al., 2018). Although attentional modulation was not included in their study, it was consistent with our observations, in that the visual rhythm was faster than the frontal rhythm by roughly 1 Hz and propagated to the anterior cortex. However, there was a mismatch in the propagation delay in the human visual to frontal cortex, which was tenfold slower compared to our observation in mice or in monkeys (Gregoriou et al., 2009). The leading of visual theta might reflect the bottom-up relay of sensory information, and the systematic reduction of the delay is functionally relevant to its purpose of faster relaying of sensory information. The delay is interpreted as a designated temporal shift for the effective relay of information, transmitting a presynaptic spike at the peak depolarization phases of postsynaptic neurons (Nowak and Bullier, 1997; Gregoriou et al., 2009). Metaphorically speaking, to selectively improve the traffic in one direction, it may be optimal for the green lights of successive crossways to be slightly delayed considering the amount of time it takes to drive through, rather than operating with zero-lag synchrony.

Then what is the generation mechanism of frontal theta observed here? Considering that our task requires an effort to withhold impulsive actions, frontal theta might originate from the interaction of the cortico-basal ganglia circuit (i.e., anterior-cingulate/medial-frontal and subthalamic nucleus) related to the conflict-related decision process (Cavanagh et al., 2011; Cavanagh and Frank, 2014; Cohen, 2014). Or the frontal theta may reflect communication between hippocampal structures and the frontal cortex (Jones and Wilson, 2005; Siapas et al., 2005), as the task required the subjects to maintain visuospatial information (i.e., direction of motion). Regarding the role of frontal theta oscillations in sustained attention, a large and consistent body of literature has discussed the top-down role of frontal theta in cognitive control (Cavanagh and Frank, 2014) and sustained attention (Clayton et al., 2015). All these works included task-engaged mental processes (e.g., keeping one’s goal in mind, overriding the prepotent response, monitoring the goal-related sensory cues). Our results provide further evidence for this top-down role of frontal theta oscillations, as we found that frontal theta increased during sustained attention in mice. Importantly, phase synchrony between frontal and visual theta increased in high-performance trials in a critical manner. Thus, frontal theta appears to be behaviorally relevant as its top-down control mode. It should be noted that the substantial work conducted in the frontal cortex has found gamma oscillations to play an important role in pro-cognitive functions (Cho et al., 2015; Kim et al., 2016). On the neuronal spike levels, the spiking activities of the frontal cortex in the high gamma band were locked to theta oscillations in the frontal and posterior parietal cortices during sustained attention (Sellers et al., 2016). In our study, we found effective modulation of low gamma, but not high gamma, by frontal theta using cross-frequency coupling analysis. Continued work with neuronal recording or optogenetic interrogation will be needed to clarify the specific role of frontal theta on neuronal activities during cognitive processing.

The counteracting behavior of visual theta compared to frontal theta was counterintuitive, but it was consistently and significantly observed across animals and days. This phenomenon of “attentional reduction” in visual theta is in agreement with recent findings in monkeys (van Kerkoerle et al., 2014; Spyropoulos et al., 2018) and ferrets (Stitt et al., 2018). In humans, it has been reported that alpha activities (8–12 Hz) in the occipital cortex reflect functional inhibition of task-irrelevant cortical areas (Jensen and Mazaheri, 2010). To the best of our knowledge, wakeful relaxation related alpha activities have not been reported in mice unlike other cross-species frequency-preserved rhythms (Buzsaki et al., 2013). In addition, compared to baseline activity, visual theta in our study increased during stimulus presentation (Fig. 2E), whereas alpha usually decreases (Fries et al., 2008), which is similar to the finding of Spyropoulos *et al.* (Spyropoulos et al., 2018). Therefore, it seems more plausible to regard the oscillatory activities observed in this study as visual theta rather than occipital alpha. However, the functional significance of such attentional reduction in the cortical microcircuit remains unknown.

What is the role of theta rhythmicity in the visual system? Recently, it has been suggested that the theta rhythmicity in the perceptual domain reflects the periodic sampling process of the sensory system (i.e., rhythmic sampling (VanRullen, 2016, 2018)). In this framework, attention periodically fluctuates around the theta-band to sample sensory stimuli, resulting in theta-rhythmic fluctuations of relevant neural activities and behavioral responses (Busch and VanRullen, 2010; Fiebelkorn et al., 2018; Kienitz et al., 2018). Still, the suppression of visual theta by attention is not explained, particularly when we consider that theta rhythms reflect the attentional sampling process. According to Spyropoulos *et al.*, the attentional reduction of visual theta was suggested to reflect “more continuous processing” of attended stimuli, and a shift to the faster rhythm was predicted (Spyropoulos et al., 2018). Moreover, Harris *et al.* reported that the phase modulation of visual theta on visual detection performance was more effective for a non-attended than attended target in humans (Harris et al., 2018). These results suggest that visual theta might be in charge of processing outside the attentional spotlight rather than involved in neural processing in the focal attentional field.

While a growing number of studies have revealed the modulation effects of visual theta on neural excitability, the neural basis for the generation of visual theta remains unknown (see 34). Recently, a predominant feedforward directionality of visual theta was found in monkeys, from lower to higher levels of the visual cortex, and to the temporal cortex (Spyropoulos et al., 2018). In addition, a colocalization of stimulus-induced gamma oscillations and visual theta oscillations was revealed (van Kerkoerle et al., 2014). Considering that the feedforward directionality of gamma oscillations in the primary sensory cortex is relatively well known, it is believed that visual theta reflects bottom-up sensory processing rather than top-down influences from higher-order visual areas. Moreover, simultaneous recording of the visual thalamic area and the posterior-parietal cortex in the ferret brain also demonstrated a strong feedforward directionality in the theta frequency band (Stitt et al., 2018), but it is not clear whether it is generated from the thalamo-cortical network or instead is locally driven.

If visual theta propagates in a feedforward direction, how can it be modulated by top-down processes such as attention? One recent study using ferret thalamus recordings suggests an answer to this question. It found that spiking activities in the pulvinar are modulated by attention, and these neurons signal in the theta frequency band during sustained attention (Yu et al., 2018). This area exhibits bidirectional anatomical connections with higher-order visual cortices (Jones, 2001), and even with the prefrontal cortex (Romanski et al., 1997). As there has also been a report that unit activities in the primate visual area V4 are modulated by the external theta driver in the prefrontal cortex (Liebe et al., 2012), it seems highly plausible that the top-down modulation of theta is generated via a cortico-thalamic input, which in turn leads to a change of thalamo-cortical connectivity. Indeed, attentional modulation of neuronal activities in the visual cortices has attracted a wide range of interest among cognitive scientists. Considering the genetic techniques uniquely applicable to the mouse model, future studies may take advantage of mouse models to understand the functional architecture and underlying circuitry of visual theta rhythmicity in association with its attentional modulation.

## METHODS

All procedures were approved by the Institutional Animal Care and Use Committees at the corresponding author’s primary affiliation and complied with NIH guidelines. Detailed experimental procedures were as follows.

### Animal Preparation

Male C57BL/6 mice were used in the experiments. The animal care and surgery were performed as described in *Extended Data - Materials and Methods*.

### Go/No-Go Task

After mice learned to discriminate the direction of the random-dot motion on a screen, their task performance was tested by randomly presenting rightward (“Go”) or downward (“No-Go”) motion of random dots that stopped at licking or after 4 s when licking was not detected. Each mouse performed 10 experimental sessions on successive days. The design of the stimulus, training, and task are described in *Extended Data - Materials and Methods*.

### Rating Task Performance

The ongoing task performance of mice was assessed by the ratio of correct answers (Hit or correct rejection) in 10 successive trials, as described in *Extended Data - Materials and Methods*.

### Electroencephalography

EEG responses were recorded at a 2-kHz sampling rate with microscrew electrodes implanted on the skull above the frontal and visual cortices. Ground and reference electrodes were implanted on the occipital bone, as described in *Extended Data - Materials and Methods*.

### EEG Analysis

Before analysis, the data were high-pass filtered at 1 Hz and then segmented from 2 s before stimulus onset to 2 s after the cessation of the stimulus. Theta oscillations were investigated by a bandpass filter at 4–12 Hz. For the time-frequency analysis, a fast Fourier transform at a 1-Hz interval with a 512-ms sliding Hanning window with a step size of 100 ms was performed. To eliminate the effect of early evoked potentials, the first 500 ms of each epoch was excluded from the main analyses. Possible cross-frequency coupling between theta and gamma oscillations was studied using the modulation index described by Tort *et al.* (Tort et al., 2010). The modulation index for phase frequency in the theta band and that for amplitude frequency in the fast frequency band (20–40 Hz for low gamma, 80–160 Hz for high gamma) were calculated. Further information on data analysis can be found in *Extended Data - Materials and Methods*. Also, several Matlab functions used in this study are available online, including calculation of spectrogram, phase locking value, instantaneous frequency difference, and time lag from given EEG signal (https://github.com/EEG-processing-scripts/matlab_functions_for_EEG).

### Statistical Testing

Non-parametric versions of the t-test (two-sided), one-way ANOVA, and circular centrality test with *α* = 0.05 were performed for statistical tests. For the correlation analysis, Pearson’s method was used with *α* = 0.05. Sample size was not previously estimated for the experiments, and blinding or randomization was not needed in this work, as the design was decided by subjects’ performance.

## Extended Data Figures

### This PDF file includes

- Extended Data - Materials and Methods

- Extended Data - Figures 1-1, Figure 1-2, Figure 2-1, Figure 3-1, and Figure 4-1.

## Extended Data - Materials and Methods

### Animals and surgery

Five male C57BL/6 mice (4–8 weeks old at time of surgery, body weight *M* = 26.1 g, *SD* = 4.7 g) were used, maintained under a 12:12 h light:dark cycle (lights on at 8:00 a.m.) in a temperature- and humidity-controlled environment. Food was available *ad libitum*, and water was mildly restricted (1 mL per day, additional water available as reward from task) from two weeks after surgery. Health status of the animals was assessed daily according to a systematic assessment protocol (Guo et al., 2014). All procedures were conducted in compliance with the Institutional Animal Care and Use Committee of corresponding author’s primary, conforming to the NIH Guide for Care and Use of Laboratory Animals (NIH Publication No. 86-23, revised 1985).

For stereotaxic surgery, mice were anesthetized with a ketamine and xylazine cocktail (120 and 6 mg/kg, respectively) by intraperitoneal injection and positioned on a stereotaxic apparatus (Model 957, Kopf Instruments, CA, USA). After shaving the head, an incision was made to expose the skull and implant sterilized screw (Asia Bolt, South Korea) electrodes above the frontal (anterior-posterior, 1.9-2.1 mm; medial-lateral, ±2.1 mm from bregma) and visual cortices (anterior-posterior, −3.7 mm; medial-lateral, ±3.7 mm from bregma). For selection of recording sites, we referred to a previous study which targeted the mouse prelimbic area to record theta oscillations using surface electrodes *in vivo* (Sauer et al., 2015), and our pilot study using high-density surface EEG array (Choi et al., 2010) which showed that the selected mouse visual area exhibited most prominent motion-specific oscillatory responses to random-dot motion stimuli. Reference and ground electrodes were implanted on the occipital bone (Paxinos and Franklin, 2004). Dental cement (Vertex Self-Curing, Vertex Dental, Netherlands) was applied over the electrodes for fixation and two polycarbonate nuts (inner diameter 3 mm, Nippon Chemi-Con, Japan) were attached to the caudal edges of the cement, for use as anchors of the custom-built head-fixation apparatus. After surgery, mice were treated with antibiotics and analgesics.

### Stimuli and apparatus

All experiments were conducted in a lightproof Faraday cage. Mice were restrained in a custom-built acrylic tube (3.5 cm in diameter) and head fixed by the polycarbonate nuts. For a reward, a water drop (3 μL) was released through a pressure equalizer tube (2 μm in diameter) with a programmable syringe pump (Fusion 200, Chemyx, TX, USA). Licking behavior was counted with a custom-built lickometer made of stainless steel wire (3 μm in diameter) and a data acquisition system (cDAQ-9147, National Instruments, TX, USA). For punishment, a single puff of air was delivered to the animal’s cheek (air pressure = 15–20 p.s.i., duration = 0.2 s), supplied with a compressed air tank and controlled by a solenoid valve. A schematic diagram of the experimental setup is shown in Figure 1a.

For visual stimuli, a random-dot motion was generated and presented through Matlab (Mathworks, Natick, MA, USA). The dot motion was composed of 100 small dots (full coherence, size = 0.5 ° in radius, field size = 60 ° in diameter, speed = 42 °/s, dot life = 1.67 ± 0.33 s from a Gaussian distribution, orientation = 90 ° or 180 °) and presented on a LCD screen (LG E1910PM-SN, 19 inches, 1280×1024 resolution), located 15 cm away from the animal’s eyes. As for the validity of using random-dot motions in the mouse model, previous studies have shown rodents are able to discriminate the direction of the random-dot motion (Douglas et al., 2006) and exhibit brain oscillatory responses similar to those found in humans (Han et al., 2017).

### Training procedure

The behavioral training procedure was consisted of three stages: (1) habituation (7 d), (2) response shaping (2–3 d), and (3) conditioning (2–4 d). The habituation stage includes animal handling (10–15 min/d) and provides animals the opportunity to adapt to the head-fixation setup inside the experimental apparatus (30 min/d) without behavioral requirements. The response shaping stage was aimed to teach animals that water can be accessed by licking the tube. During this period, a water reward was given for every lick (lick port sensor time-out = 20 s). Once the animal started to robustly lick the tube, they advanced to the conditioning stage. The conditioning stage trained animals to associate a visual stimulus with a water reward. Each trial began with a monotone sound cue (10 kHz, 70 dB, 100 ms) and a white central fixation cross (size = 3 °). After 2–3 s of the cue, a rightward moving-dots stimulus (S+, ‘Go’-motion) was presented on the screen, which subsequently turned black for 30 s for the water-tube licking. Each stimulus did not last longer than 2 min, and 100 trials were given in one day. Our pilot study of reaction time (RT), validated through video recording, showed that intentional licking results in a typical RT distribution with a peak near 1 s, whereas spontaneous licking results in a scattered RT distribution over a 2 min time period. Also, we found that licking rates decreased over time, potentially due to thirst satiation. Thus, we considered the conditioning was successfully established when the RT of the first 80 trials reached a typical RT distribution. Overall, the percentage of mice licking in response to the visual stimulus on the last day of the conditioning stage was roughly 70%.

### Go/No-Go task

In the Go/No-Go sessions (10 d, consecutive), the animals were required to discriminate the direction of the moving dots (Fig. 1B). Each trial began with a monotone sound cue (10 kHz, 70 dB, 100 ms) and a white central fixation cross (size = 3 °). After 2–3 s of the cue, the stimulus was presented for 4 s with either a rightward- (S+, ‘Go’-motion) or downward- (S-, ‘No-Go’-motion) motion which was terminated when licking was detected. The direction of the motion of each trial was randomly determined (50%) with a restriction that the same stimulus was not presented more than 4 times in a row. A licking response during the S+, which was counted as a Hit, resulted in instant reward. If the mouse did not take the reward within 20 s, the water in the tube was suctioned out. On the other hand, a licking response to the S-was counted as a FA, resulting in instant punishment and a 20 s timeout. Trials without licking responses were counted as a Miss or CR for the S+ or S-, respectively. To guide the animals to react more consciously, rather than reflexively, the response window started 0.5 s after the onset of motion (Fig. 1B). Inter-trial intervals (ITI, black screen) were 5–8 s long. Any spontaneous licking during the ITI or cue period before the response window reset the trial starting from the ITI.

The number of trials per session was determined by the animal’s performance that day (287 trials/d on average). Each session was terminated once the animal stopped licking for 30 successive trials, which we regarded as a lack of motivation for the task, and were excluded from the EEG analysis. We unexpectedly observed an obsessive licking behavior, during which the animal would indiscriminately lick the tube regardless of stimulus type. If animals licked for more than 30 trials in a row, those trials were excluded from EEG analysis. Overall, around 41% of trials were excluded.

### Behavioral state rating

As previous human and animal studies have shown, performance in cognitive tasks varies over time depending on the behavioral state, for instance due to an attentional lapse (Liebe et al., 2012; Esterman et al., 2013). To investigate the relationship between cortical theta activities and sustained attention, it was important to infer correctly the behavioral state, as either attentive or inattentive. This is particularly important in Go/No-Go tasks because mind wandering for a short period of time (i.e., attentional lapse) can be counted as a correct No-Go response. To rate each trial according to attentional state, the consistency of performance over time was analyzed.

Because the span of attentional state in humans was reported to range between 2–7 min (Esterman et al., 2013) or exceed 10 min (Clayton et al., 2015), we assumed that short-term attentional state in mouse would be similar or shorter than that of human. Then, we assumed the level of attentional state is reflected through performance in the previous trials (in the order of a few minutes). Finally, we identified fluctuations in performance over 10 successive trials by calculating the probability of correct answers (*p*_*c*_; Hit and CR), ranging from 0 (0 correct trials) to 1 (10 correct trials) with the chance level 0.5. *p*_*c*_ of *k*-th trial was defined as follows,

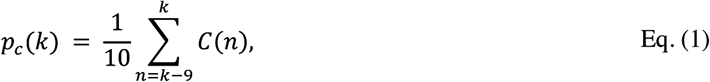

where the binary function, *C*(*n*), indicates the outcome of the trial as correct (1) or not (0). In our paradigm, 10 trials lasted approximately 2–5 min. We subsequently classified the trials into attentive state (*p*_*c*_ larger than the chance level) or inattentive state (*p*_*c*_ equal to or smaller than the chance level).

### Statistical analysis

Non-parametric versions of t-test (Wilcoxon signed-rank and rank-sum test), ANOVA (Analysis of variance; Kruskal-Wallis test), and circular-median test using CircStat Toolbox (Berens, 2009) were performed (α =.05, two-tailed). For correlation analysis, Pearson’s method was used with the same alpha level.

### EEG data acquisition and processing

EEG data were collected through Grass 8-17C amplifiers (Grass Technologies, RI, USA) with 60 Hz notch filters (gain = 50,000) and were digitized using a data acquisition system (cDAQ-9147). After band-pass filtering (1–200 Hz), the EEG data were processed on a trial-by-trial basis. To eliminate hemisphere-specific activities, EEG signals from bilateral electrodes were averaged in the time domain. To minimize the effect of early evoked activities (e.g., event-related potentials), only the EEG activities from 0.5 to 4 s after motion onset were used for statistical testing of oscillation amplitudes. We computed the mean and two-tailed 99% confidence intervals of the distribution of the overall amplitude spectrum, and the values lying outside of these confidence intervals were excluded. A time-frequency amplitude spectrogram was obtained by applying fast Fourier transform with a sliding Hanning window (bin size = 512 ms, step size = 100 ms) and by taking absolute real parts from the epochs, which resulted in a 1 Hz frequency resolution. For the time-frequency spectrogram visualization, the amplitudes of the pre-stimulus periods (2 to 0.5 s before stimulus onset) were averaged and subtracted in a frequency-wise manner.

### Cross-frequency coupling between theta and gamma oscillations

To investigate cross-frequency coupling between theta and gamma oscillations, we calculated the modulation index (MI) as described in Tort *et al.* (Tort et al., 2010). In brief: (i) raw EEG epochs were band passed (phase frequency: 4–12 Hz, amplitude frequencies: 6 to 200 Hz with 2.5 Hz-step sliding window and 5 Hz width) with 5^th^ order Butterworth filters. (ii) The time series of the phase of theta oscillations denoted as 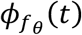, and the time series of the amplitude of the gamma oscillations denoted as 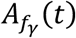, were extracted via Hilbert transform. (iii) The phase of the theta oscillations was divided into *N* bins from -*π* to *π*. In each bin, the average of 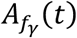 was calculated and then normalized by dividing the sum over the *N* bins and was denoted as *P*_*j*_ for the *j*-th bin. Here, *P*_*j*_ equals the probability density function of the gamma amplitude distribution over the theta phase ranging from 0 to 1. (iv) An entropy index, *H*, was calculated by the definition of Shannon entropy, given by

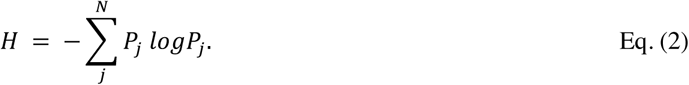

If the gamma oscillations are independent from theta phase, *P*_*j*_ is 1/G; therefore *H* = log (*N*). (V)The statistical deviance of the measured distribution (*P*) from the uniform distribution was calculated by adopting Kullback-Leibler distance by the following formula,

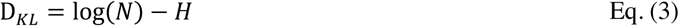

where D_*KL*_ stands for the Kullback-Leibler distance. (vi) Finally, MI was defined with a *Z*-score of observed entropy as the following formula indicates.

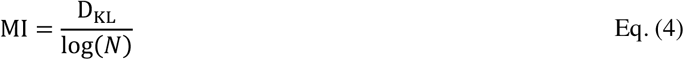

In the current study, we set *N* equal to 30 (i.e., bin size = 12°).

### Generalized Linear Model analysis of CFC

Single-trial GLM analyses were performed to investigate the increased level of theta-gamma coupling during attentive state is modulated by either theta amplitude or attentive state, or both. The model was composed of following equation,

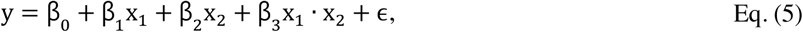

where y, x_1_, x_2_, and ϵ represent MI, theta amplitude, attentional state (inattentive = 0, attentive = 1), and error term, respectively. The term x_1_ · x_2_ represents a two-way interaction between theta amplitude and attentional state. GLM fitting is done by a Matlab built-in function (`fitlm.m`).

### Phase synchrony analysis of theta activities

To quantify the interaction between theta oscillations in the frontal and visual areas, phase synchrony was estimated by joint phase histograms and by calculating phase locking values (PLV) (Lachaux et al., 1999). To obtain joint phase histograms, a Matlab built-in function for two-dimensional histogram (`hist2.m`) was used. To calculate PLV, first, instantaneous phase of the theta oscillations was calculated via Hilbert transform and was denoted as *ϕ* _Fr_(*t*) and *ϕ*_Vis_(*t*) for the frontal and visual areas, respectively. Second, PLV was calculated in each stimulation period excluding the first 0.5 s (i.e., from *t* = 0.5 to *t* = 4) as follows,

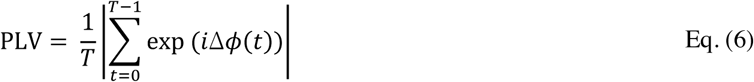

where Δ*ϕ* (*t*) = *ϕ* _Fr_(*t*) − *ϕ*_Vis_(*t*). PLV ranges from 0 (i.e., absence of phase locking) and 1 (i.e., perfect phase locking).

A cross-correlation function *ρ*_*X,Y*_(Δ*t*) of visual and frontal theta oscillations, X(*t*) and *Y*(*t*), respectively, was calculated in each trial excluding the first 0.5 s (i.e., from *t* = 0.5 to *t* = 4), according to

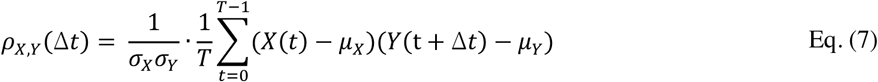

in which *μ* and *σ* indicate the mean and standard deviation. The time delay, *τ*, was obtained from;

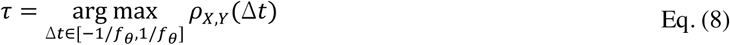

where arg max indicates the arguments of the maxima (i.e., the points that the cross-correlation functions are maximized). Positive values of *τ* indicate the temporal lead of visual theta over frontal theta and vice versa. The resolution of *τ* was 0.5 ms (2,000 Hz sampling).

## Extended Data - Figures

**Fig. 1-1.**
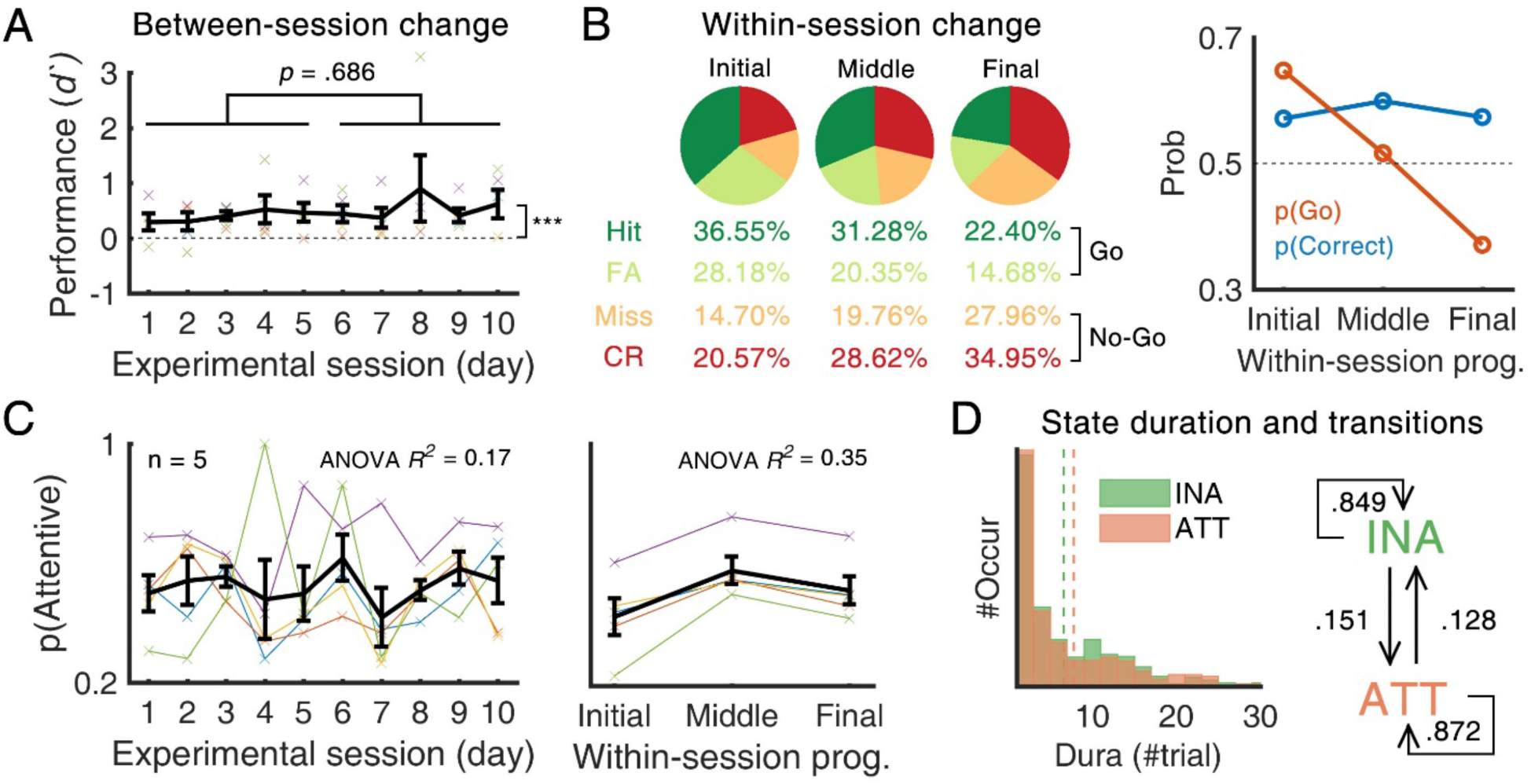
Performances changing across- and within-session. **(A)** Performance varies across sessions. Across the 10 days of sessions, overall behavioral performance (*d’, M* = 0.475, *SE* = 0.08) was significantly larger than chance level, *Z* = 5.845, *p* < .001. Learning effect, calculated by the difference of mean *d’* between first five days (*M* = 0.399, *SE* = 0.07) and last five days (*M* = 0.551, *SE* = 0.21), was not significant, *Z* = 0.405, *p* = .686. Error bars represent ± 1 SEM. (**B**) Performance varies within a session. Splitting trials of single session into three different groups, initial (first 1/3), middle (second 1/3), and final (last 1/3), change of behavioral performance within session was analyzed. In general, proportion of Go response trials (*i.e*., Hit and FA) was higher at initial phase, and gradually decreased, suggesting the change of animals’ motivational level to water reward. Interestingly, despite the decrease of probability Go response, p(Go), overall probability of correct (*i.e*., Hit and CR), p(Correct), showed relatively small variation within session. (**C**) Behavioral state changes across- and within-session. Changes of proportion of ‘attentive’-labeled trials (*p*_*c*_ > 0.5) as a function of experimental session (left) and as a function of within-session progress (right) were analyzed. To calculate the effect size, one-way ANOVA (Kruskal-Wallis test) was performed. We found experimental session explained 17% of the variance, while the within-session progress explained 35% of the variance, suggesting the larger effect size of motivation-related factors relative to the learning-related factors. Error bars represent ± 1 *SEM*. (**D**) Duration and transitions of behavioral state. Behavioral state showed transitions (i.e., transition from inattentive to attentive, and vice versa) and stationary phase with certain duration (inattentive: 6.60 trials in average, attentive: 7.81 trials in average) (left). Using Markov-chain model with two-state process, this state transition was summarized in a state diagram (right).

**Fig. 1-2.**
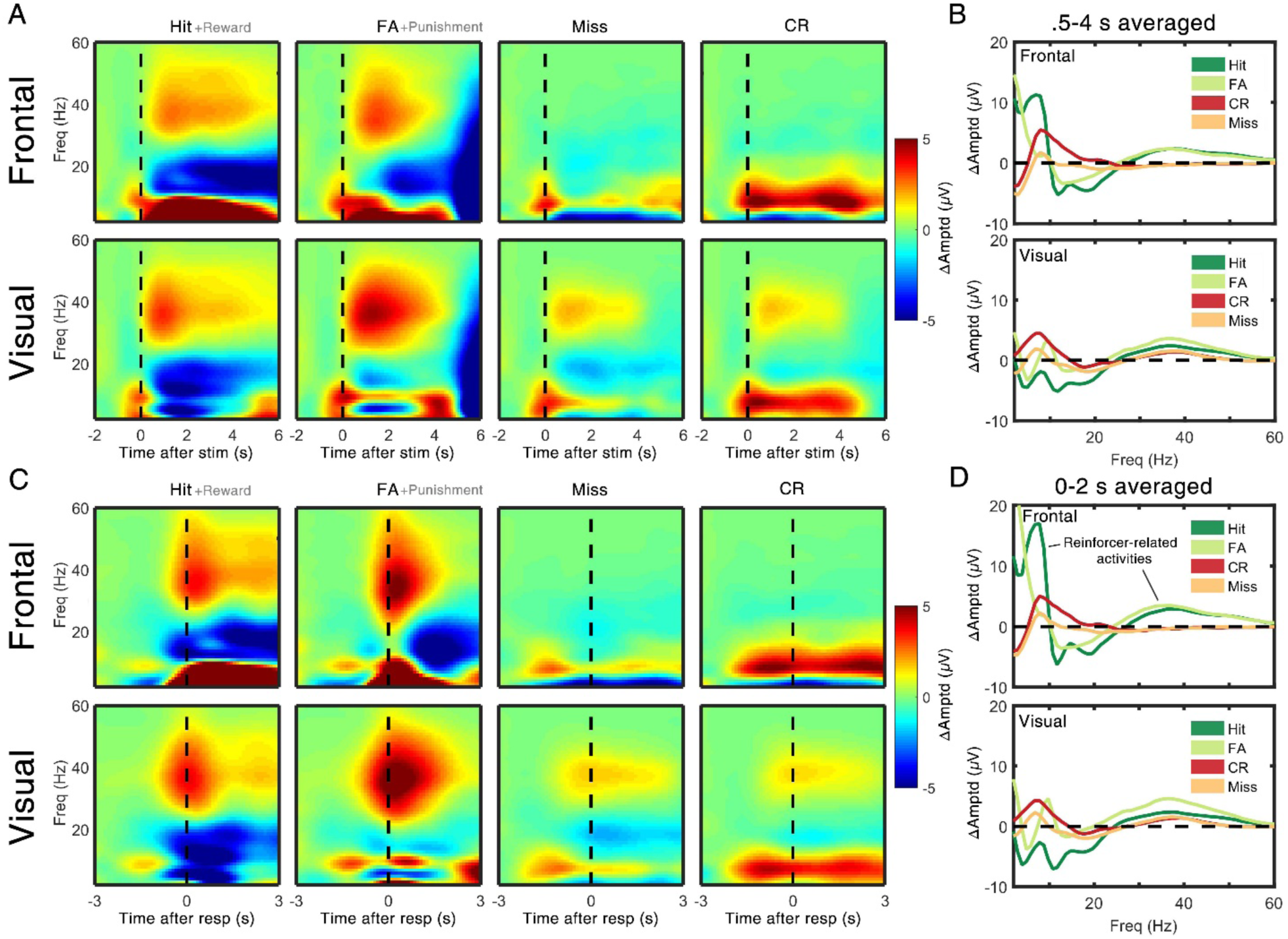
Grand-averaged EEG responses during Go/No-Go task. **(A)** Stimulus-locked grand-averaged amplitude spectrogram of four behavioral outcomes. Black dashed lines indicate the onset of visual stimulus. Hit and FA trials exhibited relatively large changes in oscillatory amplitudes, compared to those of Miss and CR trials. These large activities were considered as reinforcer-related activities (i.e., water reward, air puff punishment, motor movement, etc.) as they were locked to the moment of reaction (See panel c). **(B)** Averaged (t = 0.5 – 4 s after motion onset) amplitude spectrum of spectrograms in (a). **(C)** Same as (a), but response-locked. For the trials which do not contain behavioral response (CR, Miss), mean RTs of the session from Hit and FA trials were used to align the epoch. **(D)** Same as (b), but response-locked.

**Fig. 2-1.**
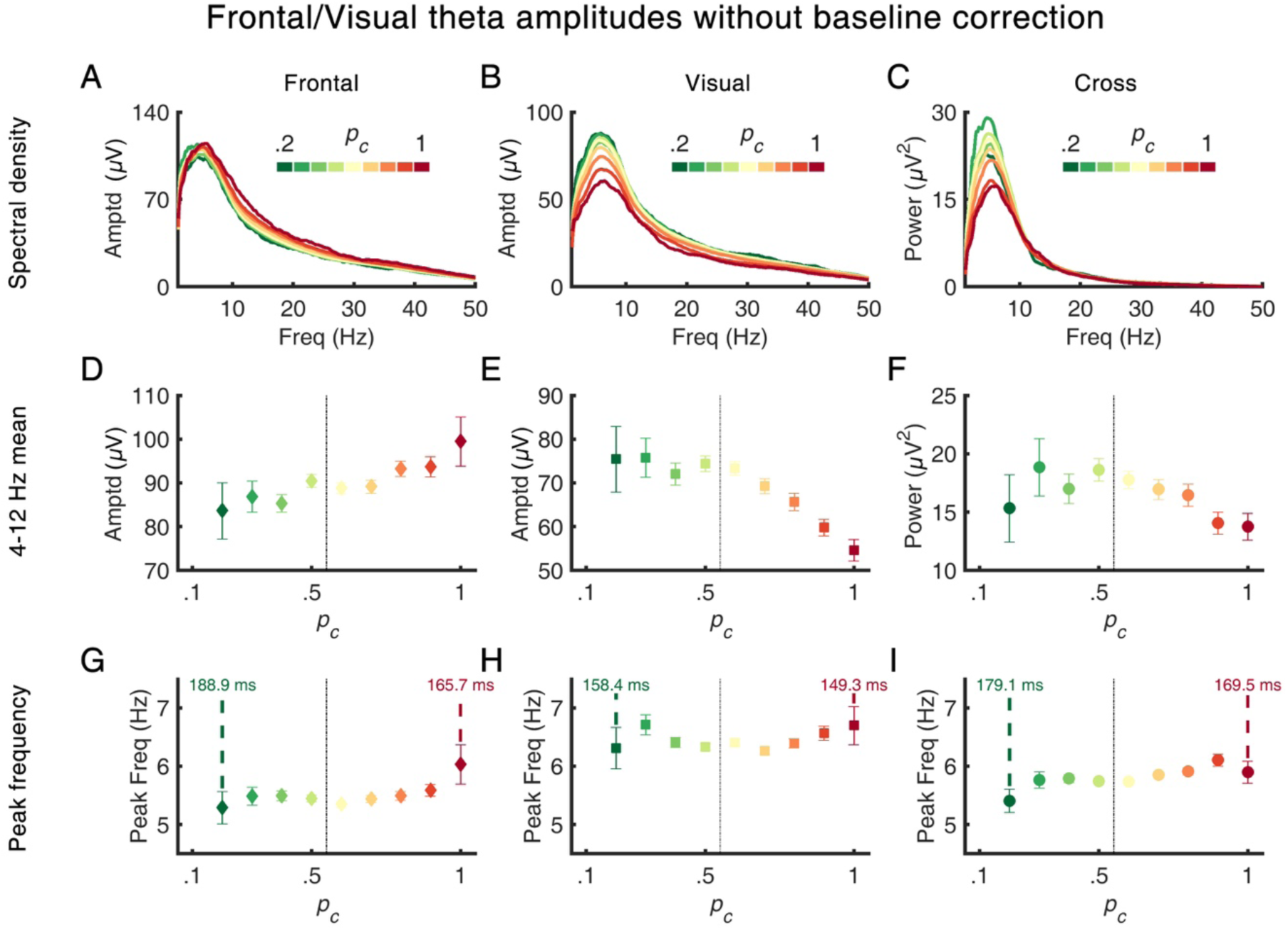
Baseline-uncorrected amplitude- and cross-spectral densities. **(A)** Mean amplitude- or cross-spectrum of CR trials (n = 2,411, t = 0.5–4 s) calculated from the frontal. Error bars represent ± 1 *SEM*. **(B)** Same as (**A**), from the visual. **(C)** Same as (**A**), from cross-spectrum of the frontal and visual. **(D)** Mean frontal theta (4–12 Hz) amplitude. Error bars represent ± 1 *SEM*. **(E)** Same as (**D**), from the visual. **(F)** Same as (**D**), from cross-spectrum of the frontal and visual. **(G)** Mean peak frequency of the frontal theta amplitude spectrum. Peak values were calculated in trial-by-trial basis. Colored numbers illustrate the duration of one cycle of each frequency. Error bars represent ± 1 *SEM*. **(H)** Same as (**G**), from the visual. **(I)** Same as (**G**), from cross-spectrum of the frontal and visual.

**Fig. 3-1.**
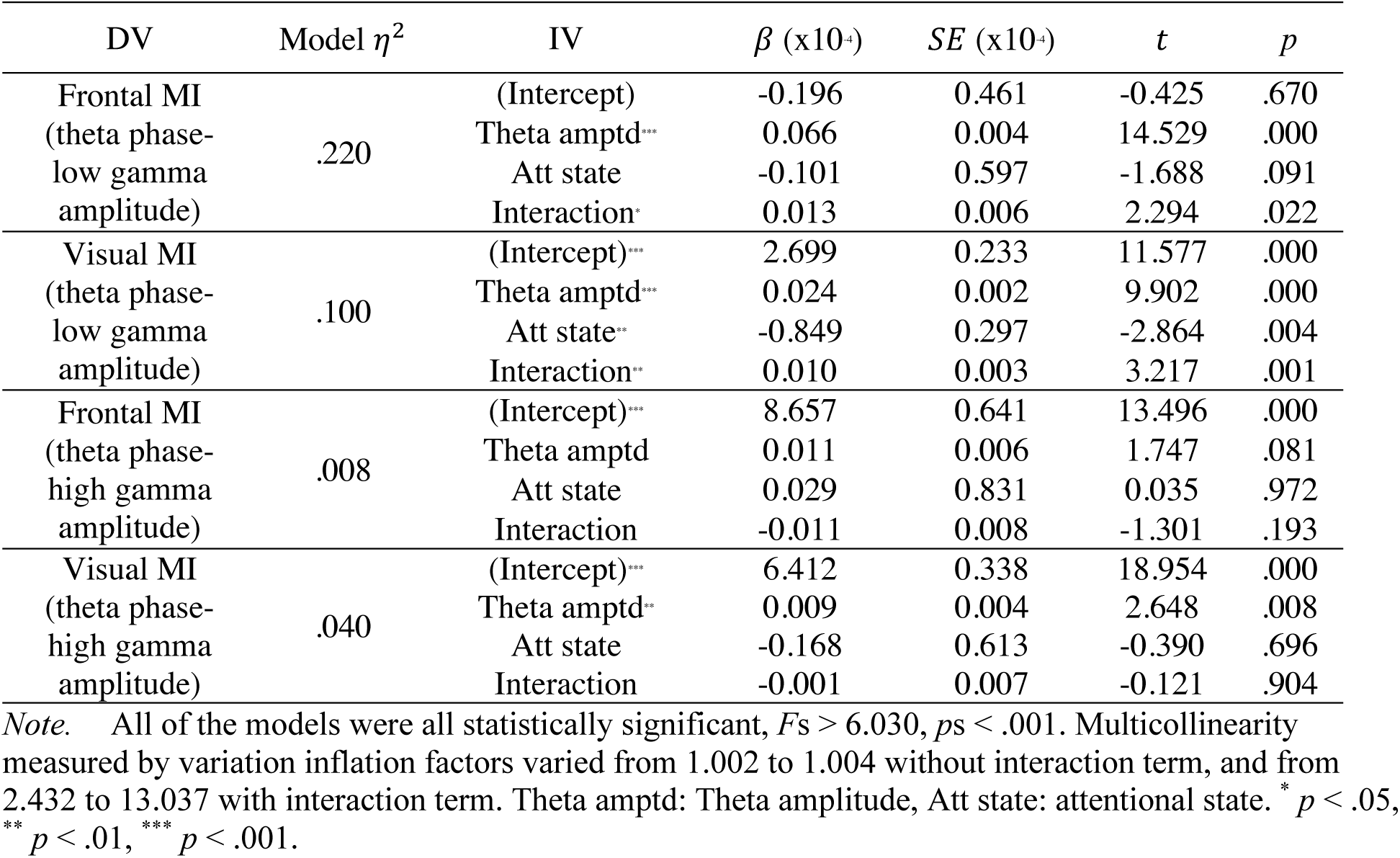
GLM analysis of cross-frequency coupling. To dissect the confounding effect of theta and attention on MI, MIs were further analyzed through Generalized Linear Model (GLM) with three predictors: theta amplitude, attentive state, and two-way interaction of theta amplitude and attentive state (See Extended Data - Materials and Methods for details). As a result, we found statistically significant main effect of theta amplitude on theta-low gamma coupling in both regions (frontal: *t* = 14.529, *p* < .001, visual: *t* = 9.902, *p* < .001) and on theta-high gamma coupling of visual (*t* = 2.648, *p* < .01). The effect of theta amplitude on frontal theta-high gamma MI did not reach to statistical significance (*t* = 1.747, *p =* .081). Also, the effect of attentional state on MI was statistically significant only for visual theta-low gamma coupling (*t* = −2.864, *p* < .01) with non-significant effect on the other couplings (|*t|*s < 1.688, *p*s > .091). Most importantly, the two-way interaction between theta amplitude and attentional state was statistically significant for theta-low gamma coupling in both regions (*t*s > 2.294, *p*s < .05), but not for theta-high gamma coupling (|*t*|s < 1.301, *p*s > .193). Such two-way interaction with positive slope value (*ß* = 0.010 × 10^−4^ for frontal, *ß* = 0.013 × 10^−4^ for visual) suggests the effect of theta amplitude on theta-gamma coupling gets stronger with attentive state, especially for theta-low gamma coupling in both regions. In short, the change of CFC observed in this study can be best explained by a synergetic effect of theta amplitude and attentional state, rather than by single factor.

**Fig. 4-1.**
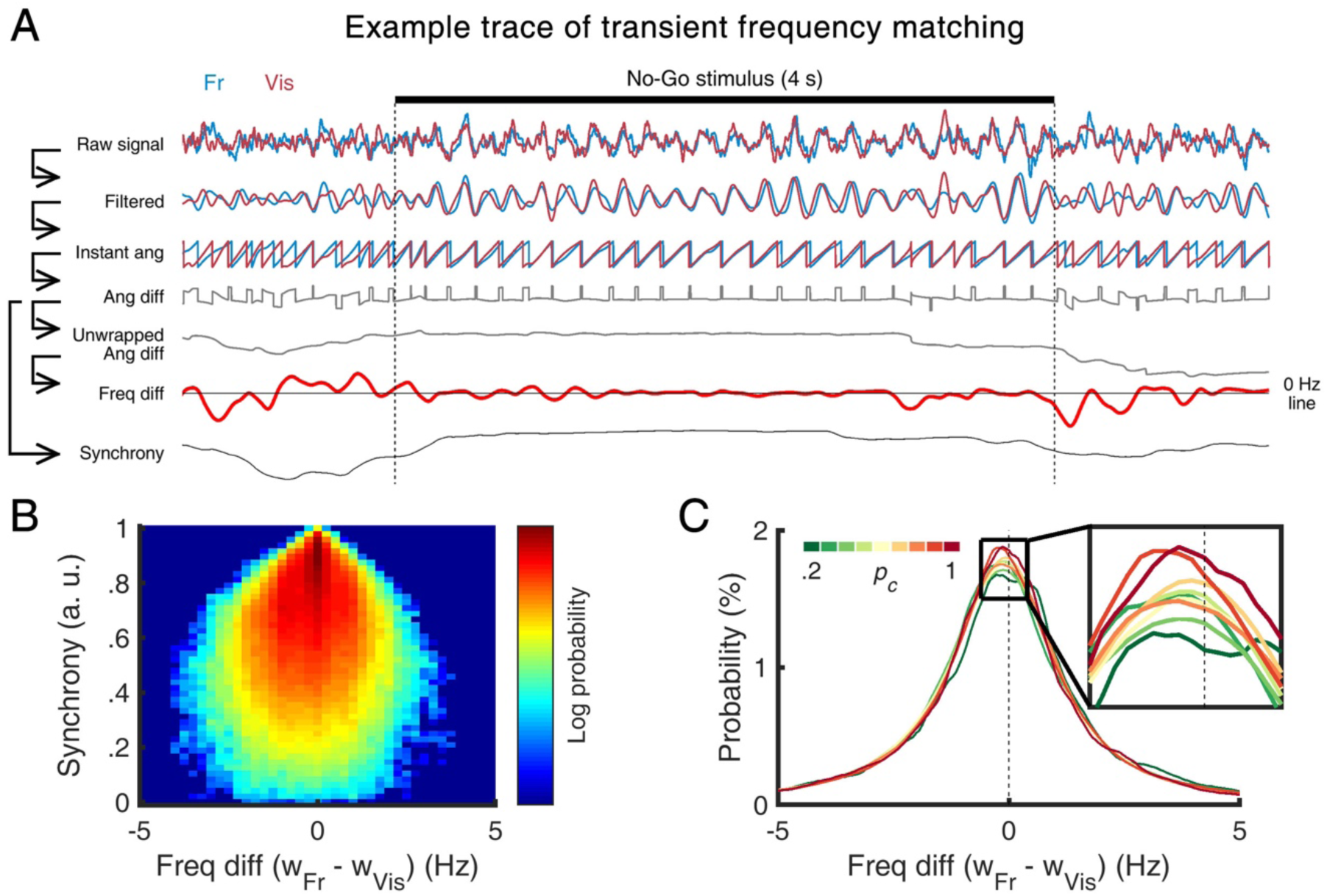
Transient frequency matching during fronto-visual theta synchrony. (**A**). Example single-trial data showing transient frequency matching and increased phase synchrony. To calculate frequency difference and phase synchrony, raw data were converted into Hilbert instantaneous angle after band-pass filtering. Frequency difference was obtained by taking derivate of unwrapped angle difference between two channels’ data. Fr: Frontal, Vis: Visual, Ang: Angle, Diff: difference, Freq: frequency. **(B)** Two-dimensional histogram of instantaneous frequency difference and phase synchrony during the No-Go period of all trials (irrespective of attentional state). High synchrony was observed when frequency difference is low. Note that synchrony value was calculated with 1 s width and 10 ms sliding temporal window, and instantaneous frequency difference was time-averaged with same size sliding window. W_Fr_: Instantaneous frequency of frontal theta, W_Fr_: Instantaneous frequency of visual theta. **(C)** Histogram of instantaneous frequency difference, drawn separately for each *p*_*c*_ group. Probability of having transient frequency matching was higher during the attentive than during the inattentive state.

